# An Ultralong Bovine CDRH3 that Targets a Conserved, Cryptic Epitope on SARS-CoV and SARS-CoV-2

**DOI:** 10.1101/2022.04.06.487306

**Authors:** Matthew J. Burke, James N.F. Scott, Thomas Minshull, Peter G. Stockley, Antonio N. Calabrese, Joan Boyes

**Affiliations:** School of Molecular and Cellular Biology, Faculty of Biological Sciences, University of Leeds, Leeds, LS2 9JT, UK; Astbury Centre for Structural Molecular Biology, Faculty of Biological Sciences, University of Leeds

## Abstract

The ability of broadly neutralising antibodies to target conserved epitopes gives them huge potential as antibody-based therapeutics, particularly in the face of constant viral antigen evolution. Certain bovine antibodies are highly adept at binding conserved, glycosylated epitopes, courtesy of their ultralong complementarity determining region (CDR)H3. Here, we used a SARS-naïve, bovine ultralong CDRH3 library and mammalian cell display, to isolate a bovine paratope that engages the SARS-CoV and SARS-CoV-2 receptor-binding domain (RBD). This neutralises viruses pseudo-typed with SARS-CoV Spike protein but not by competition with RBD binding to ACE2. Instead, using differential hydrogen-deuterium exchange mass spectrometry and site-directed mutagenesis, we demonstrate that this ultralong CDRH3 recognises a rarely identified, conserved, cryptic epitope that overlaps the target of pan-sarbecovirus antibodies (7D6/6D6). The epitope is glycan-shielded and becomes accessible only transiently via inter-domain movements. This represents the first bovine anti-sarbecovirus paratope and highlights the power of this approach in identifying novel tools to combat emerging pathogens.

## Introduction

The severe-acute respiratory syndrome coronavirus (SARS-CoV)-2 pandemic has highlighted the immense threat that our societies face from highly transmissible, zoonotic pathogens, especially when there is negligible pre-existing immunity. The occurrence of three coronavirus spill-over events in the last 20 years, together with the huge reservoirs of these viruses in species such as bats, indicates that coronaviruses will continue to threaten the population globally. Perhaps more worryingly, the transfer of other pathogens has also been occurring at an accelerated rate. Although vaccines haven proven remarkably successful at slowing SARS-CoV-2 transmission and preventing severe disease [1-3], this will not be the case for every pathogen. Moreover, not all individuals mount an effective immune response to vaccines, meaning that it is vital to develop complementary therapeutics that can be rapidly deployed in response to a new pathogen.

Monoclonal antibodies (mAbs) provide alternative treatments, particularly for individuals with weakened immune systems [4] as well as for the general population in the absence of an effective vaccine. Numerous virus-neutralising antibodies (nAbs) against the SARS-CoV-2 homotrimeric Spike glycoprotein have been isolated, which can potently reduce infection and confer protection *in vivo*. Most nAbs target the Spike receptor binding domain (RBD), which promotes cell entry via interactions with the human angiotensin-converting enzyme (hACE2) [5-7]. Unfortunately, the majority of these nAb are vulnerable to escape mutations [8], including many mAb-based therapies that have previously entered the clinic under emergency use authorizations [9]. Given that many other RBD mutations can be tolerated without compromising Spike expression or Spike-hACE2 interactions [10], antigenic escape from mAb therapies was anticipated. This was recently realised by the Omicron variant, which has 15 mutations in the RBD and is resistant to neutralisation by most currently available mAb therapeutics [11, 12].

The ideal mAb therapeutic would therefore target a conserved neutralising epitope to minimise its vulnerability to escape mutations, whilst also increasing its breadth of activity against related viruses. Various human Abs have been identified that cross-react among sarbecoviruses and which neutralise via distinct mechanisms [13-17]. For example, AB-3467, sterically blocks ACE2/RBD interaction [18, 19] whereas S2H97 appears to convert the Spike protein to a post-fusion state [15]. It remains possible, however, that additional conserved epitopes will be identified using different or non-conventional antigen-binding structures. Neutralising antibodies against such epitopes may provide broader protection, not only for SARS-CoV and SARS-CoV-2 but also the inevitable future spill-over events of other zoonotic sarbecoviruses.

Bovine ultralong antibodies provide a promising new toolset. Previous studies demonstrated that broadly neutralising anti-HIV-1 Abs could be isolated from a cow immunised with a stabilised HIV-1 Env [20]. The highest affinity antibody (NC-Cow-1) engages with the CD4 binding site of Env and potently neutralises a large panel of HIV-1 variants [20]. The remarkable breadth of this broadly neutralising bovine Ab can be attributed to its unconventional paratope structure that allows it to target a small footprint on Env and reduce its vulnerability to escape mutations [21]. Similarly broad and potent ultralong nAbs have been isolated against Foot-and-mouth disease virus from infected cattle [22]. Crucially, the paratope from NC-Cow-1 was successfully transferred to a human antibody scaffold with minimal loss of activity, establishing the feasibility of generating humanised bovine ultralong antibodies as therapeutic tools [23].

The structure of bovine ultralong antibodies makes them exceptionally well suited to achieve broad and potent neutralisation [24-26]. This subset of Abs possesses a long CDRH3 of 40-71 amino acids (aa) which forms an extended β-strand stalk supporting a disulphide-bonded ‘knob’ domain [25, 27, 28]. The latter is entirely responsible for all direct antigen interactions and by sitting on top of a long β-stranded stalk, it can punch through glycan coats to reach normally occluded epitopes [20, 21]. The entire bovine ultralong CDRH3 repertoire is encoded by recombination of the same three gene segments: V_H1-7,_ D_H8-2_ and J_H2-4_, each of which has a defined role in generating and stabilising its structure [25, 26, 29] and which lends itself well to the specific isolation of ultralong CDRH3 sequences (Fig. 1a).

**Figure 1:**
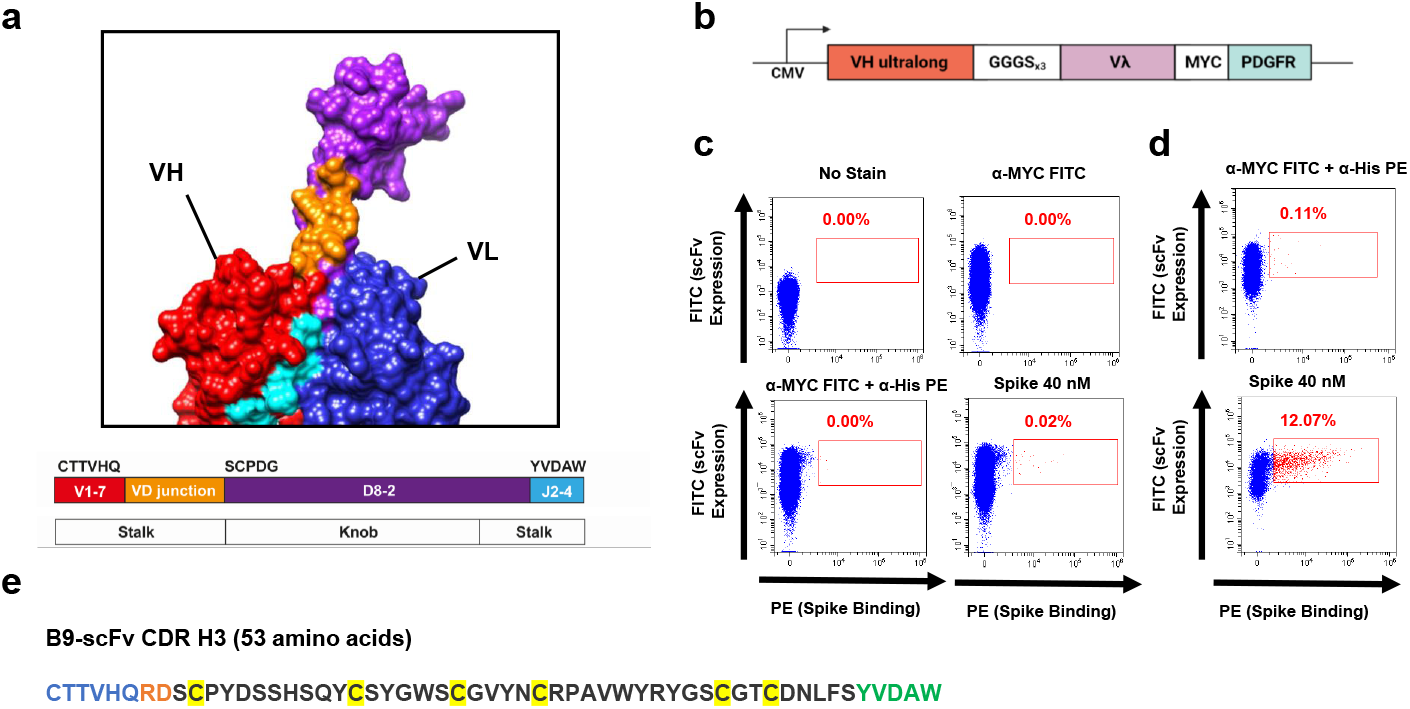
Cell display and binding of ultralong scFvs to SARS-CoV-2 Spike protein. **a**) Ultralong CDRH3 sequences are generated by recombination of the same three gene segments: V_H1-7,_ D_H8-2_ and J_H2-4._ These contribute to specific parts of the solvent exposed CDRH3 stalk and knob domain, as indicated by the colour-coded gene segments and CDRH3 structure. **b)** Construct to express membrane-bound scFvs with an ultralong V_H_. **c)** Upper: FACS plots without (left) and with (right) incubation with α-Myc-FITC. Lower: FACS plots showing α-His-PE and α-Myc-FITC staining in the absence (left) or presence (right) of 40 nM Spike. The red box shows cells expressing scFvs that bind Spike. **d)** scFvs that bind to Spike are markedly enriched after two rounds of plasmid-based selection. 293T cells were transfected with Round 2 plasmid library incubated without (upper) and with (lower) 40 nM Spike. All samples were stained with α-His-PE and α-Myc-FITC (1:100). **e)** The amino acid sequence of B9-scFv. The regions encoded by V_1-7_ (blue), the VD junction (orange), D_8-2_ (dark grey) and J_2-4_ (green) are shown. Cysteine residues are highlighted in yellow.

Here, we sought to isolate ultralong bovine heavy chains with neutralising activity against SARS-CoV-2 and related coronaviruses. We capitalised on previous findings that these pair with a relatively invariable Vλ light chain [25, 29], to generate a single chain variable fragment (scFv) scaffold into which ultralong heavy chain-only libraries can be cloned and expressed. Using mammalian cell surface display and His-tagged S2P SARS-CoV-2 Spike glycoprotein, we isolated an ultralong scFv (B9-scFv) from a SARS-CoV-2-naïve heavy-chain library that binds to SARS-CoV-2 RBD, all current SARS-CoV-2 variants and, with a ≈50x stronger interaction, to SARS-CoV RBD. B9-scFv does not compete with ACE2 binding but neutralises SARS-CoV pseudo-typed lentiviruses with an IC_50_ of 468 nM, likely by destabilising the prefusion complex. Consistent with this, the epitope localises to a cryptic cleft on the inner face of the RBD that becomes available only transiently by inter-domain movements. This epitope has been identified only once previously as the target for two other pan-sarbecovirus antibodies [30] when multiple, repeated immunisations were required. Remarkably, here we isolate a bovine broadly reactive CDRH3 from a library of <1 × 10^4^ sequences. This attests to the huge potential of the bovine system as a source of broadly reactive antibodies that can protect against emerging pathogens and their variants.

## Results

### Cell surface display of bovine ultralong scFvs

To establish a cell-surface display platform for the isolation of ultralong CDRH3s, we generated a library of ultralong bovine paratopes by amplifying variable exons from the leukocyte genomic DNA (gDNA) of two adult cows. Ultralong CDRH3s were enriched by nested PCR and gel excision, resulting in an initial heavy chain library with >96% ultralong CDRH3s (Supplementary Fig. 1a). The purified amplicons were inserted into the pBovShow cassette (Fig. 1b) that resulted in them being joined to an invariant Vλ light chain (Vλ-LC) via a flexible linker. Ultralong scFvs were expressed following transient transfection into 293T cells, as determined by flow cytometry (Fig. 1c). Notably, cell surface expression of over 70% of scFv clones was achieved (Supplementary Fig. 1b), suggesting the invariant Vλ light chain pairs with most ultralong heavy chains, even in an scFv format.

### Isolation of a bovine ultralong scFv that binds to the SARS-CoV-2 Spike glycoprotein

Next, we screened our ultralong scFv library for binding to the recombinant full-length SARS-CoV-2 S-2P glycoprotein [31] by transient transfection into 293Ts and subsequent flow cytometry. Initially, a small population of cells expressing scFvs that bind Spike protein were isolated (0.02%; Fig. 1c); these were enriched by two further rounds of plasmid recovery, re-transfection and flow cytometry (Fig. 1d), resulting in a sharp increase in the proportion of Spike-binding scFvs (Fig. 1d). Sequencing the scFv library at each stage of enrichment allowed the original library diversity to be estimated at <1 × 10^4^ unique sequences (Supplementary Fig. 1c). Given this low level of initial diversity, it is remarkable that scFvs that bind SARS-CoV-2 Spike protein are present.

To isolate these Spike-binding scFv(s), we cloned our enriched library into lentivirus (LV) vectors and generated LV particles pseudo-typed with VSV-G. These were transduced into 293T cells at a low titre to achieve few scFv sequences per cell. After puromycin selection and expansion, only 0.42% of the transduced cells bound Spike protein (Supplementary Fig. 2a, b). From this population, we isolated 15 single cell clones (SCCs) that interacted with 40 nM Spike. Three of these SCCs harboured a single scFv and remarkably, the sequence of all three scFvs was identical at the nucleotide level. This sequence, termed B9-scFv (Fig. 1e), encodes a 53 amino acid ultralong CDRH3 that interacts with the SARS-CoV-2 Spike (Supplementary Fig. 2b, lower). B9-scFv accounted for 53% of all scFvs from the LV-transduced cells after a single selection for Spike binding, and this increased to 83% upon a further round of enrichment by flow cytometry. These data therefore suggest that the ultralong B9-scFv likely accounts for much the anti-Spike activity in our library.

### The epitope for B9-scFv is within the SARS-CoV-2 RBD

To further characterise B9-scFv, we purified the SARS-CoV-2 Spike sub-domains (Fig. 2a), S1 (aa 2-682), S2 (aa 686-1211), and RBD-SD1 (aa 319-591) via IMAC (Supplementary Fig. 3a). Cells transiently expressing B9-scFv display clear binding to the S1 domain and the RBD, but not to equivalent concentrations of the S2 domain (Fig. 2b). These data therefore localise the B9-scFv epitope to RBD residues 319-591.

**Figure 2:**
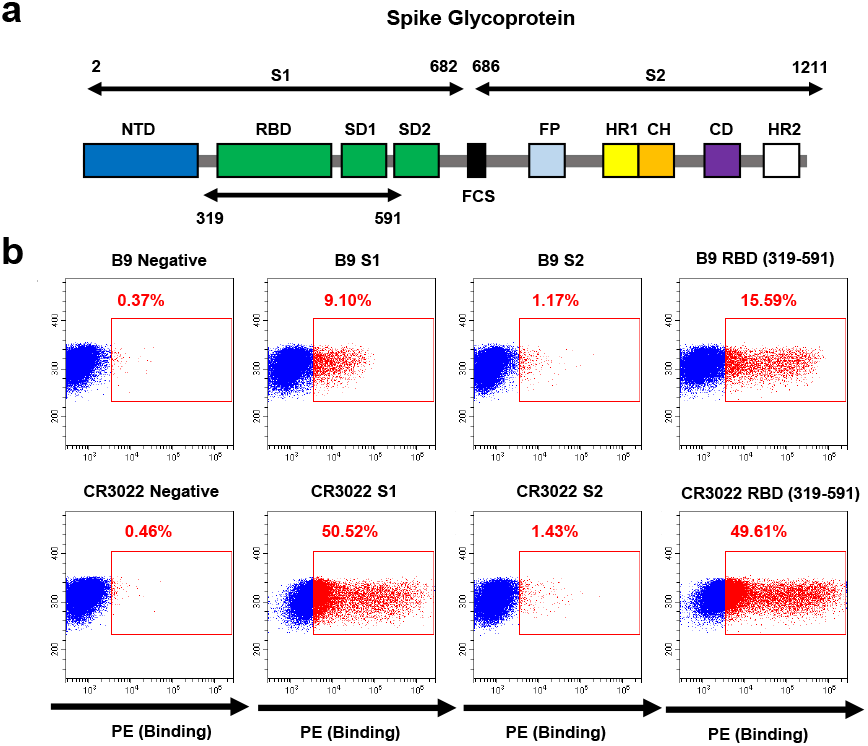
Isolation of an ultralong scFv that binds to wildtype SARS-CoV-2 Spike receptor binding domain. **a)** Cartoon depicting the SARS-CoV-2 Spike protein sub-domains across aa 2-1211. NTD: N-terminal domain, RBD: receptor binding domain, SD1: sub-domain 1, SD2: sub-domain 2, FCS: furin cleavage site, FP: fusion peptide, HR1: heptad repeat 1, CH: central helix, CD: connector domain, HR2: heptad repeat 2. **b)** FACS plots of 293T cells transfected with plasmids encoding B9-scFv (top panels) or CR3022-scFv (bottom panels). These were incubated with or without 2 μM of the 8xHis-tagged SARS-CoV-2 sub-domains shown, and 1:100 α-His-PE fluorescent antibody.

Although B9-scFv readily binds 40 nM Spike, markedly higher levels of RBD (≈2 μM) are necessary to achieve a similar effect (Supplementary Fig. 3b). Similar disparities in apparent affinity have been reported previously for the interactions of engineered ACE2-Fc constructs with RBD-monomers and full-length trimeric Spike [32]. These were attributed to differences in avidity due to the increased number of binding sites provided by the trimeric Spike compared to the RBD. While these data suggest B9-scFv has an epitope within SARS-CoV-2 RBD, they also suggest its binding may be relatively weak.

### B9-scFv is resistant to receptor binding motif (RBM) mutations

Bovine ultralong CDRH3s typically recognise conserved epitopes. If this is the case for B9-scFv, we would expect its binding will be unaffected by mutations in any of the SARS-CoV-2 variants of concern (VOC). To investigate this, the B9-scFv sequence was cloned upstream of a poly-His tag, expressed in 293Ts and purified by IMAC (Supplementary Fig. 4a). 293T cells were then transiently transfected with an expression vector for SARS-CoV-2 Spike (Wuhan-Hu-1 + D614G) and cell-surface Spike expression was confirmed by staining with a positive control scFv (CR3022-scFv), which binds to a conserved RBD epitope between SARS-CoV-2 and SARS-CoV [33] (Supplementary Fig. 4b, upper). Titration of purified B9-scFv gives concentration-dependent binding to Spike transfected cells, that is enriched compared to a negative control bovine ultralong scFv (137-scFv; Supplementary Fig. 4b). We next examined B9-scFv binding to SARS-CoV-2 Spike protein variants; crucially, binding is maintained to all of the mutations in the commonly circulating SARS-CoV-2 variants, including D614G, N501Y, E484K, Y453F, L452R and K417N from the Alpha, Beta, Gamma, Delta and Omicron variants (Fig. 3a, b). Not only does this imply B9-scFv recognises SARS-CoV-2 Spike glycoprotein in its native state, but also suggests B9-scFv may be broadly reactive.

**Figure 3:**
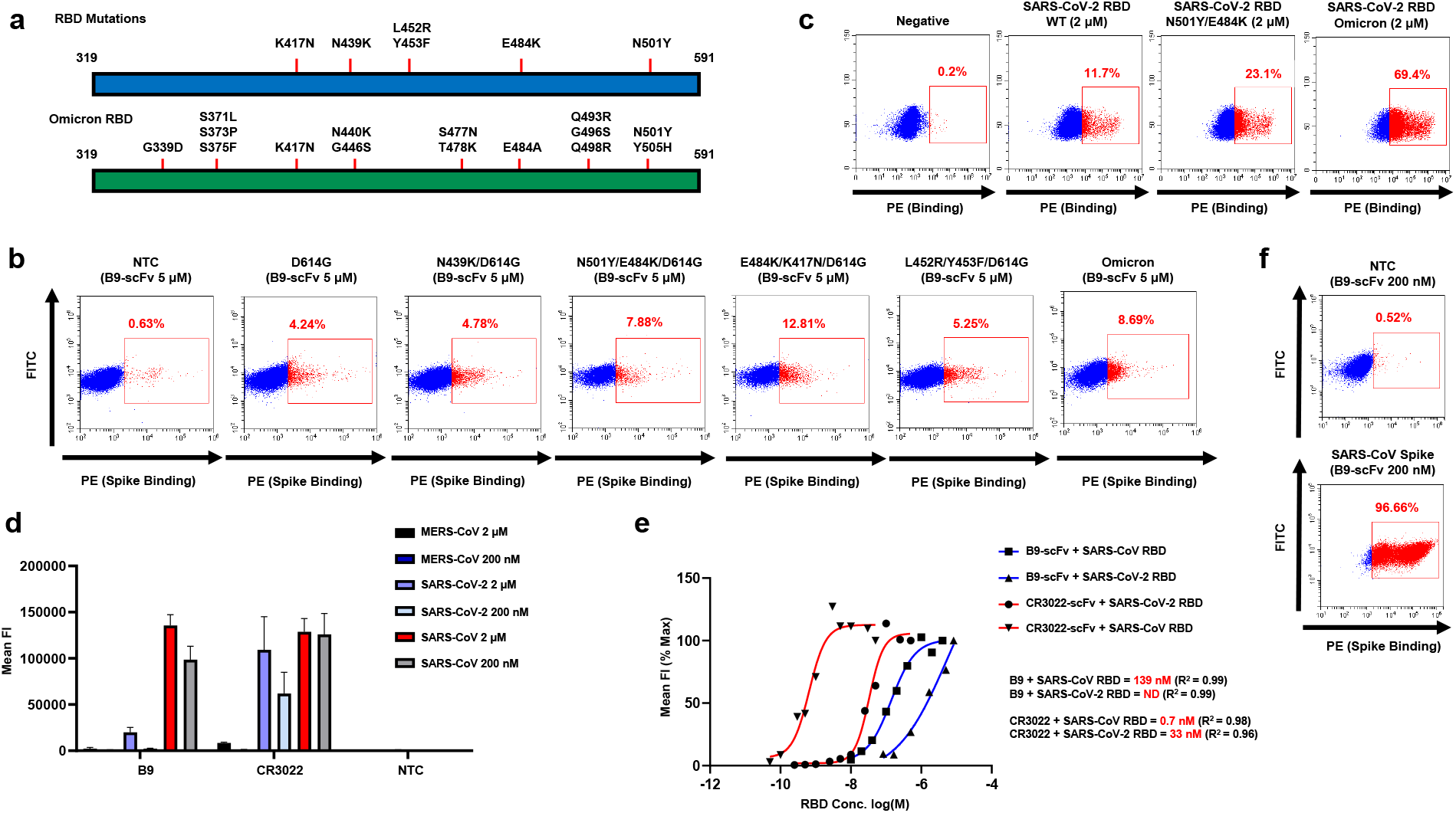
Purified B9-scFv is broadly reactive and binds strongly to SARS-CoV RBD. **a)** Cartoon depicting the positions of the mutations in the SARS-CoV-2 Spike RBD (aa 319-591) (upper) and Omicron (lower). All mutations are on the Wuhan-Hu-1 + D614G background. **b)** Non-transfected (NTC) 293T cells and cells transfected with expression vectors for the mutant Spike proteins shown were incubated with 5 μM 8xHisTagged B9-scFv. Samples were then stained with α-His-PE (1:100). **c)** Non-transfected 293T cells (NTC) and 293T cells transfected with B9-scFv incubated with the RBD variants shown at 2 μM; left: WT RBD, middle: N501Y/E484K RBD, right: Omicron RBD. RBD binding is indicated by the red box. All samples were stained with α-His-PE and α-Myc-FITC (1:100). **d)** Non-transfected 293T cells (NTC) or 293T cells transfected with B9-scFv or CR3022-scFv were incubated with 2 μM or 200 nM of purified 8xHisTagged MERS-CoV, SARS-CoV-2 or SARS-CoV RBD (Urbani variant), followed by staining with α-His-PE antibodies (1:100). Mean fluorescence intensity of PE staining was used to quantify binding. Errors bars indicate standard deviation of three independent repeats. **e)** 293T cells were transfected with B9-scFv or CR3022-scFv and incubated with titrations of purified SARS-CoV-2 or SARS-CoV RBD (Urbani variant). The apparent K_D_ of the interaction between cell surface scFv and purified RBD was approximated from the non-linear analyses of log(concentration)-response plots. Data points were plotted as a percentage of the mean fluorescent intensity obtained at the highest RBD concentration tested in each case. Estimated K_D_ and R^2^ values are indicated. **f)** Binding of purified B9-scFv (200 nM) to 293T cells that were non-transfected (upper) or transfected with full-length SARS-CoV Spike (Urbani variant; lower). All samples were stained with α-His-PE antibody (1:100).

To further confirm B9-scFv binding to the SARS-CoV-2 VOC, we purified SARS-CoV-2 RBDs (aa 319-591) harbouring various mutations and incubated these with 293T cells expressing B9-scFv on the cell surface. Consistent with the data in Fig. 3b, B9-scFv bound to RBDs carrying the mutations associated with the Alpha lineage as well as to the RBD of the hypermutated Omicron variant, with no significant loss of affinity compared to wild type Spike (Fig. 3c). Notably, these RBD mutations found in Omicron are also found in the Beta and Gamma variants. These data therefore add weight to the idea that B9-scFv binds a conserved epitope.

### B9-scFv binds to SARS-CoV with nanomolar affinity

Given its potential broad reactivity, we next sought to determine if B9-scFv binds to other coronvirus RBDs. The corresponding sequences of SARS-CoV RBD (aa 319-591) and MERS-CoV RBD (aa 368-586) were cloned, and the proteins purified from 293T cells (Supplementary Fig. 3c). Surprisingly, the interaction between cell surface expressed B9-scFv and SARS-CoV RBD was markedly stronger than that observed with equivalent amounts of SARS-CoV-2 RBD. Maximal detectable binding by FACS was observed at 2 μM SARS-CoV RBD and this was only partially reduced at 200 nM. In contrast, the interaction with SARS CoV-2 RBD was modest at 2 μM RBD and scarcely detectable at 200 nM (Fig. 3d). Both B9-scFv and the SARS-specific human CR3022-scFv were relatively unreactive with the MERS-CoV RBD at all concentrations tested (Fig. 3d), suggesting B9-scFv has specificity for SARS-CoVs.

The small difference in binding of B9-scFv to 2 μM and 200 nM SARS-CoV RBD indicates a nanomolar affinity. To further investigate this, binding of cell surface expressed B9-scFv to a range of RBD concentrations (10 nM – 4 μM) was measured (Fig. 3e), allowing an approximate K_D_ of 139 nM to be calculated. Using this same method, we approximated the K_D_ for CR3022-scFv interactions with the SARS-CoV (0.7 nM) and SARS-CoV-2 (33 nM) RBDs, values that are comparable to those reported in the literature [34]. The binding of B9-scFv to SARS-CoV-2 RBD increased almost linearly up to the maximum analysed concentration of 5 μM RBD and therefore the affinity of the interaction could not be estimated (Fig. 3e).

We next tested if purified B9-scFv recognises the SARS-CoV RBD in the context of the Spike trimer. Consistent with our previous results, B9-scFv binds more strongly to cells expressing the SARS-CoV Spike than the SARS-CoV-2 Spike, as much lower concentrations of B9-scFv (<200 nM) are needed to label cells expressing this glycoprotein (Fig. 3f). Collectively, these data suggest that B9-scFv cross-reacts with viruses in the *Sarbecovirus* subgenus and binds with much higher affinity to SARS-CoV compared to SARS-CoV-2

### B9-scFv neutralises SARS-CoV pseudo-typed viruses

Given that B9-scFv appears to be broadly reactive, based on its resistance to all current SARS-CoV-2 RBM mutations and its cross-reactivity with the SARS-CoV RBD (Fig. 3b, c, d, e, f), we next sought to determine if it also neutralises virus infectivity. We therefore capitalised on the higher affinity of B9-scFv for SARS-CoV Spike to ask whether B9-scFv neutralises pseudo-typed lentivirus particles. As can be seen in Figure 4a, B9-scFv fully neutralises lentiviral particles pseudo-typed with the SARS-CoV Spike (Urbani variant) when tested at 70 μg/ml but has no consistent effect on an equivalent titre of SARS-CoV-2 (Wu-1-D614G) pseudo-typed virus, correlating with previously observed differences in estimated affinity (Fig. 3d). Importantly, B9-scFv does not reduce the infectivity of control VSV-G pseudo-typed lentivirus at 70 μg/ml (Fig. 4a) whereas titration experiments demonstrate that it neutralises SARS-CoV pseudo-typed lentiviruses with an IC_50_ of 468 nM (Fig. 4b).

**Figure 4:**
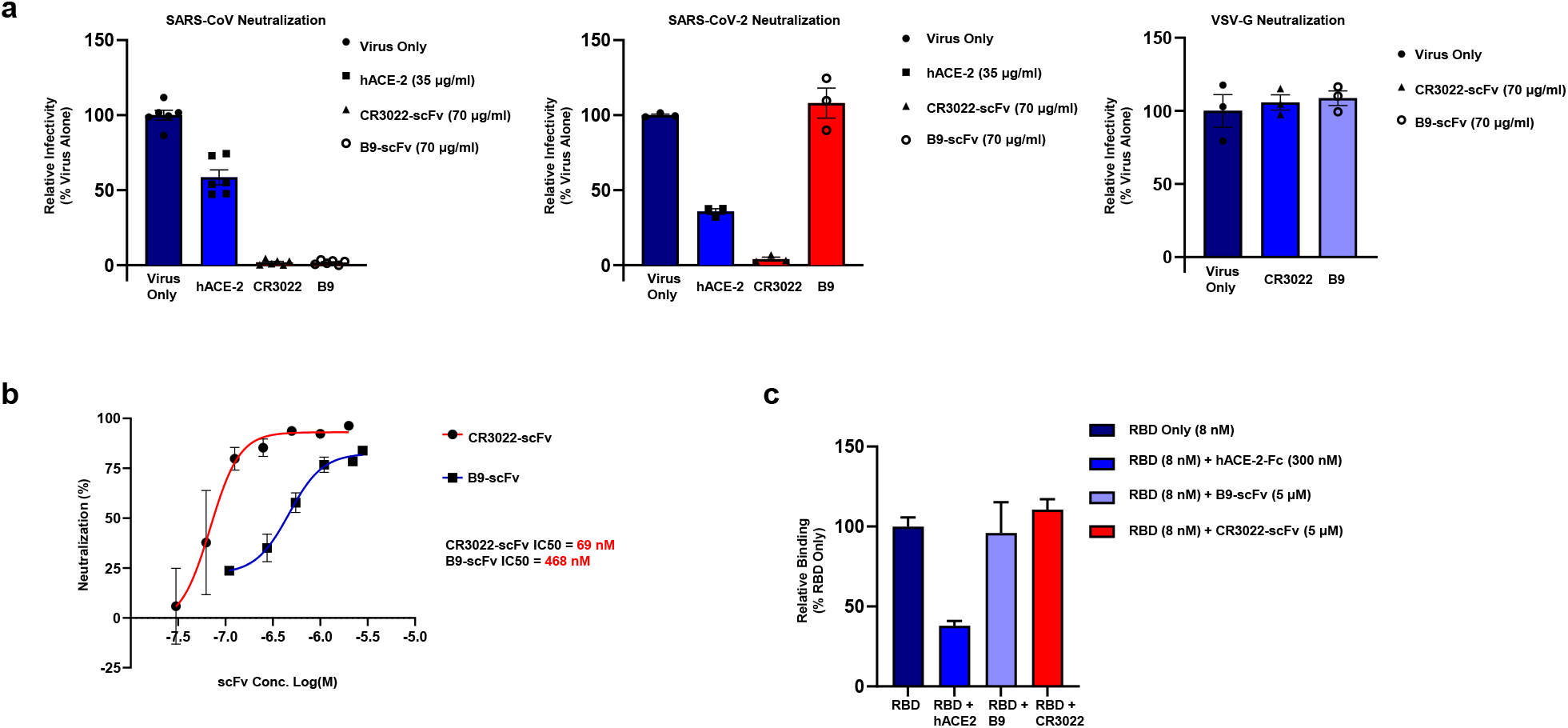
B9-scFv neutralises SARS-CoV but does not compete with ACE2. **a)** Comparable titres of lentiviruses pseudo-typed with SARS-CoV Spike protein (left), SARS-CoV-2 (Wu-1/D614G; middle) or VSV-G (right), and expressing a luciferase reporter gene, were incubated with the indicated concentrations of hACE2, CR3022-scFv or B9-scFv. 293T-hACE2 cells were transduced with the lentiviruses and the infectivity measured 48 hours later by relative luciferase units. Infectivity is compared to pseudo-typed lentivirus only in each case. The data shown is the average of three (VSV-G, SARS-CoV-2) or six (SARS-CoV) data points (±SEM). **b)** SARS-CoV pseudo-typed lentiviruses were incubated with 0.02 - 2 μM CR3022- or 0.04 - 3 μM B9-scFvs and used to transduce 293T-hACE2 cells. Neutralisation was measured by luciferase activity in the infected cells. Data comprises three independent repeats (±SEM). **c)** SARS-CoV RBD (8 nM) was incubated with the indicated amounts of ACE2-Fc (300 nM), B9-scFv (5 μM) and CR3022-scFv (5 μM) prior to measuring RBD binding to 293T-hACE2 cells via staining with α-His-PE antibody (1:100) and flow cytometry. Data comprises three independent repeats (±SEM).

In complementary experiments, we used B9-scFv in a competition-binding assay to test if it prevents purified SARS-CoV RBD from binding to hACE2-expressing cells. Although there is significant competition between soluble hACE2-Fc (300 nM) and cell surface ACE2 for RBD binding (Fig. 4c), high concentrations of B9-scFv (5 μM) or CR3022-scFv (5 μM) did not prevent SARS-CoV RBD from binding to cell surface ACE2. The distinct lack of competition in this context suggests that the mechanism of B9-scFv neutralisation is unlikely to directly involve interference with ACE2 binding.

### B9-scFv binds to a cryptic site on the RBD

Given that the Spike proteins of SARS-CoV and SARS-CoV-2 are 76% identical [35], we took advantage of the higher affinity of B9-scFv for SARS-CoV RBD to better localise its epitope. To this end, we performed differential hydrogen-deuterium exchange mass spectrometry (HDX-MS) of SARS-CoV RBD in the absence (SARS-CoV RBD only) or presence of B9-scFv (SARS-CoV RBD + B9-scFv). A 2-minute exposure to deuterium revealed two main regions of the SARS-CoV RBD with significantly reduced deuterium uptake in the presence of B9-scFv (Fig. 5a and Supplementary Fig. 5a). Protected region 1 includes three peptides involving RBD residues 449-467 and spans from β7 near the ACE2 interacting region, through the β7-β8 loop to an inner face of the RBD (Supplementary Fig. 5b). Region 1 can be further resolved to just residues 456-467 due to the identification of a shorter protected peptide (Supplementary Fig. 5a). In contrast, region 2 comprises only a single peptide of residues 551-565 within sub-domain 1 and appears to be markedly less protected than region 1 (Fig. 5a). Given that residues 542-591 can be removed with only a small reduction in B9 binding to SARS-CoV RBD (Supplementary Fig. 5c), Region 2 does not appear to be the main epitope. We therefore focused on region 1.

**Figure 5:**
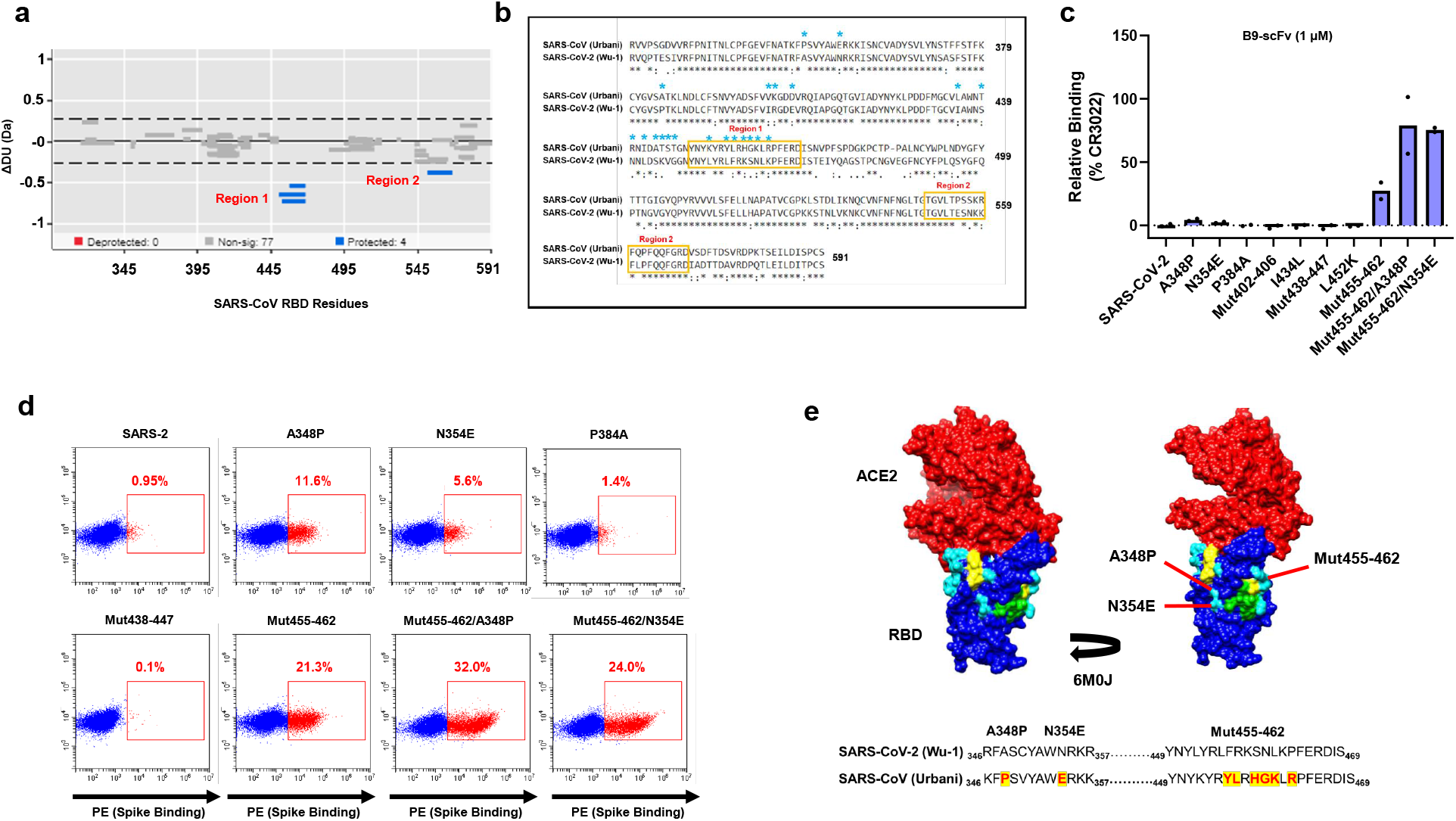
B9-scFv binds to a conserved, cryptic site on the RBD: **a)** Wood’s plots showing the summed differences in deuterium uptake in SARS-CoV RBD at two minutes of exposure to deuterium, comparing RBD alone to RBD in the presence of B9-scFv. Wood’s plots were generated using Deuteros. Peptides coloured in blue are protected from exchange in the presence of B9-scFv. Peptides with no significant difference between conditions, determined using a 99% confidence interval (dotted line), are shown in grey. For each time point and condition, three replicate measurements were performed. **b)** Amino acid sequence alignments of SARS-CoV (Urbani) and SARS-CoV-2 (Wu-1) RBDs (aa 319-591) used in this study. Orange boxes around the sequence indicate a protected region on the SARS-CoV RBD when incubated with B9-scFv as identified by HDX. A cyan asterisk above the sequence indicates the residues in SARS-CoV-2 that were mutated to their SARS-CoV equivalent in the binding studies in (**c**). **c) & d)** 293T cells were transiently transfected with constructs expressing the full-length SARS-CoV-2 (Wu-1) Spike, or the SARS-CoV-2 Spike carrying the indicated mutations. Cells were then incubated with 1 μM B9-scFv for 1 hour and stained with α-His-PE (1:100) prior to flow cytometry fluorescence analysis. Binding was quantified as the percentage of binding with a positive control CR3022-scFv (300 nM). Quantification of binding is shown in (**c**) N=2; selected FACS plots are shown in (**d**). **e)** Modelling the proposed epitope of B9-scFv onto a crystal structure of SARS-CoV-2 RBD (blue) bound to ACE2 (red) (PDB: 6M0J). HDX protected region 1 (RBD residues 449-467) is in yellow, while residues within 5 Å of this are in green or cyan. Those in cyan indicate those mutations tested in (**c & d**), arrows indicate the mutations that improve binding. The alignments of the relevant regions of the SARS-CoV and SARS-CoV-2 RBDs are presented underneath. Mutations that improve binding in **(c, d)** are highlighted, in red and labelled above.

Region 1 contains a candidate motif, _463_PFERD_467,_ that is fully conserved between SARS-CoV-2 and SARS-CoV and may explain the cross-reactivity of B9-scFv (Fig. 5b). To further investigate this, residues surrounding this motif were mutated in full-length SARS-CoV-2 Spike to their equivalents in SARS-CoV (Supplementary Fig. 5d), with the hypothesis that only mutations proximal to the *bona fide* epitope will improve binding. The various SARS-CoV-2 mutants were transfected into 293T cells and subsequently incubated with a concentration of B9-scFv (1 μM) at which binding to SARS-CoV-2 Spike is minimal. Three mutations strengthen the interaction with B9-scFv, namely A348P, N354E and Mut455-462 (Fig. 5c, d). Of these, Mut455-462 has the largest effect and is located adjacent to the _463_PFERD_467_ motif in both the primary amino acid sequence and tertiary structure of the RBD (Fig. 5c, d, e). Similarly, although the A348P and N354E mutations are distal from _463_PFERD_467_ in the primary amino acid sequence, they are in very close proximity in the tertiary structure (Fig. 5e, Supplementary Fig. 5b) and are situated on the opposite side of _463_PFERD_467_ relative to Mut455-462 (Fig. 5e). In contrast, mutations located away from _463_PFERD_467_ on the globular RBD, such as P384A, Mut402-406, Mut438-447, I434L and L452K, had no effect on B9-scFv binding to the mutated SARS-CoV-2 Spike (Fig. 5c, d, e). When combined, the A348P, N354E and Mut455-462 mutations increased binding of B9-scFv to SARS-CoV-2 to levels comparable with the positive control (CR3022-scFv; Fig. 5c). The region mapped by these experiments therefore corresponds to the smaller HDX protected peptide of residues 456-467 in region 1 (Supplementary Fig. 5a), supporting the idea that this is the epitope recognised by B9-scFv.

### Potential mechanism of neutralisation

Interestingly, in the context of a 1-up, 2-down conformation SARS-CoV Spike, the epitope for B9-scFv is relatively inaccessible on all protomers (Fig. 6a). This site has, however, previously been proposed to be transiently exposed by interdomain movements [30]. Notably, a glycan from the NTD of the adjacent protomer (N165) is thought to block access to this region by inserting itself in the volume left by the RBD when it is in the “up” conformation [36]. This glycan forms the remaining contact between NTD_B_ and RBD_A_ to stabilise RBD-up via a “lock and load” mechanism ([36]; Fig 6b). Transient domain movements that allow B9-scFv binding would break this glycan contact, potentially destabilising the Spike complex. Likewise, two previous neutralising antibodies that target this site cause destabilisation of the prefusion Spike complex [30] and shedding of the SARS-CoV-2 S1 domain.

**Figure 6:**
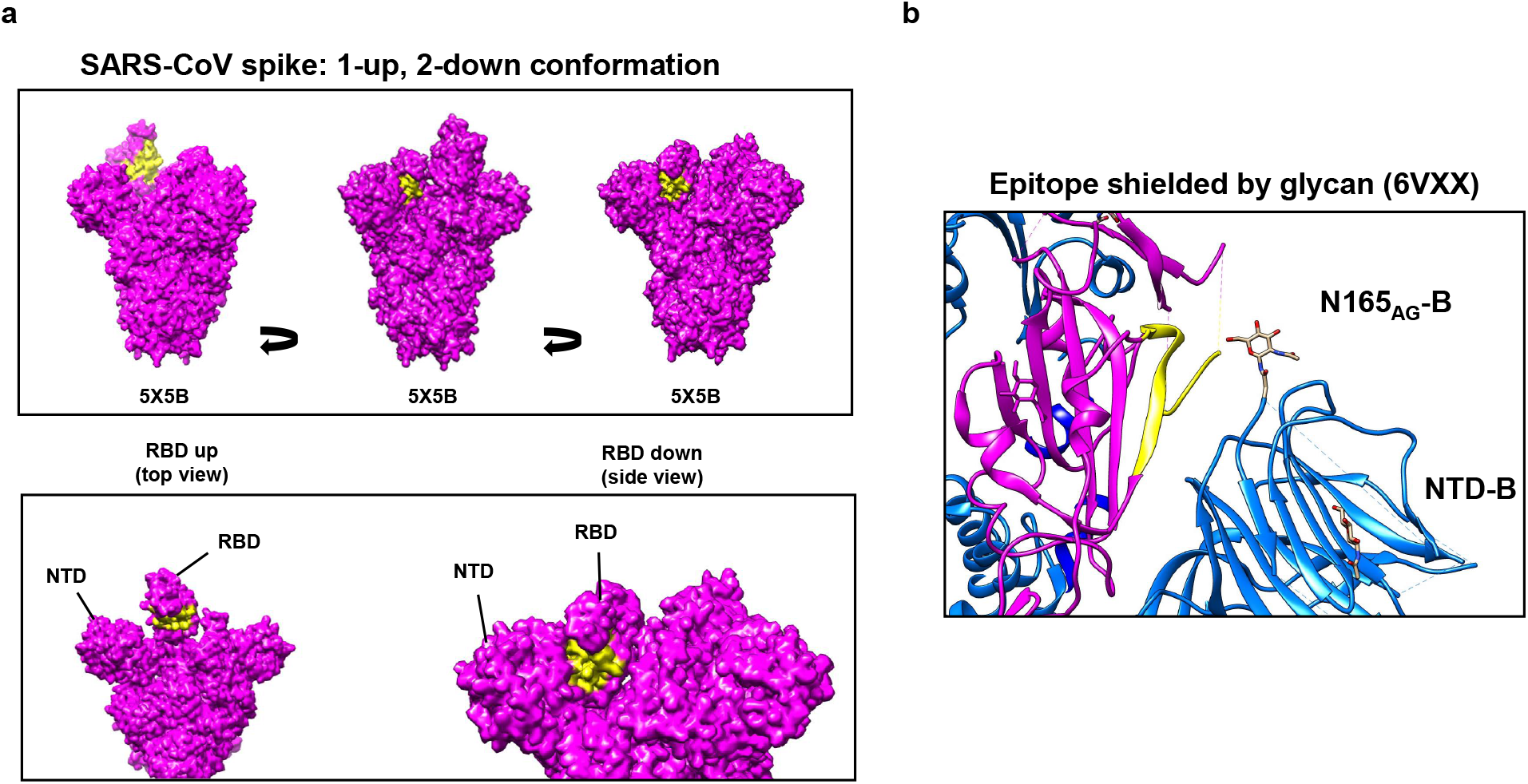
B9-scFv binding to the cryptic epitope interferes with a stabilising glycan interaction: **a)** A molecular surface map of the full-length SARS-CoV spike (PDB: 5×5B) coloured in magenta was generated in UCSF Chimera. The yellow surface indicates the proposed epitope of B9-scFv; this region is relatively occluded in all contexts. In an RBD-down this epitope is pressed against the N-terminal domain of the neighbouring protomer, while in the RBD-up conformation it is likely inaccessible without extensive clashes. **b)** Ribbon diagram showing the epitope for B9-scFv in yellow on Spike protomer A (magenta). This lies adjacent to protomer B (blue) where a glycan on N165 potentially occludes conserved epitope. PDB file: 6VXX.

Notably, B9-scFv engages this region with knob domain only binding, as disruption of the ultralong knob domain by mutagenesis abolishes the interaction (Supplementary Fig. 5e). Taken together, these data provide strong evidence that B9-scFv binds at a site largely made up of conserved residues in the β7-β8 loop and causes destabilisation of the Spike trimer. The increased strength of its interaction with SARS-CoV can largely be accounted for by a handful of residues that vary between SARS-CoV-2 (Wu-1-D614G) and SARS-CoV (Urbani).

## Discussion

The increasing frequency of zoonotic transfers, together with the possibility of rapid viral evolution, highlights the urgent need for broadly active therapeutics to fight emerging pathogens. Using our novel scFv display system, which specifically expresses bovine ultralong heavy chains in mammalian cells, we have isolated a bovine ultralong scFv (B9) that engages the receptor binding domain (RBD) of the SARS-CoV-2 and SARS-CoV Spikes at a conserved but cryptic site. Only two neutralising antibodies targeting this site have previously been isolated (7D6/6D6) and they were shown to induce destabilisation of the prefusion Spike complex [30]. Consistent with this, B9-scFv neutralises SARS-CoV pseudo-typed lentiviral particles (IC_50_ 468 nM) but not by competing with RBD binding to ACE2. Instead, it likely causes destabilisation of the prefusion Spike complex in part by interfering with a critical glycan contact. It is notable that isolation of the 7D6/6D6 antibodies required five immunizations of mice with either SARS-CoV-2 S-2P or a combination of SARS-CoV-2 Spike, SARS-CoV Spike and MERS-RBD; by contrast, our present study isolated B9-scFv from a small library of bovine CDRH3s. In fact, the identification of an ultralong CDRH3 targeting a seemingly rare neutralising epitope is made even more remarkable by the limited size of our initial library (<1 × 10^4^ unique sequences). Overall, this study clearly demonstrates how our novel approach can be used to rapidly isolate ultralong CDRH3s that target conserved sites of vulnerability on viral antigens.

Bovine ultralong antibodies possess the longest known CDRH3 regions. It is likely that the extended β-stranded stalk helps to punch through glycan coats on some viral antigens to reach occluded, functionally conserved epitopes, while it has been shown that the disulphide-bonded loops can engage a target using a compact surface area. Together, these structural features increase the resistance of ultralong neutralising antibodies to viral escape mutations [21]. In keeping with previously studied ultralong Abs, B9 engages its epitope with knob domain-only binding since disruption of the knob by mutation completely abrogates its interaction with SARS-CoV RBD (Supplementary Fig. 5e). Unusually, however, B9-scFv has a truncated ascending and descending β-stranded stalk [25, 26], with fewer residues at the VD junction (_101_RD_102_) and no alternating tyrosine motif at the 3’ end of D8-2. These features will likely impact the length, angle, and flexibility of the stalk and may influence how the disulphide-bonded loops of the B9 knob domain engage the RBD.

Notably, when the epitope for B9-scFv was mapped onto a model of a trimeric SARS-CoV Spike in the 1-up, 2-down RBD conformation, it appears relatively inaccessible in every context. Our HDX and mutagenesis data show B9-scFv’s epitope shares several key contacts with the previously identified cross-neutralising antibodies 7D6 and 6D6, including N/E354 on β1 and residues 457-467 on the β7-β8 loop of the RBD. This implies that our scFv is also likely to clash with the N-terminal domain (NTD) from the adjacent protomer in the context of the trimeric Spike (Fig. 6c). The adjacent NTD also contains a crucial aminoglycan moiety at N165. While this has been shown to be critical for gating of the RBD opening, it may also act to shield this conserved region from antibody recognition (Fig. 6b) [30, 36]. Indeed, previous models suggest this cryptic epitope is only made accessible by transient movements of the RBD and NTD, and subsequent antibody binding acts to destabilise the trimer. This ultimately leads to neutralisation by the destruction of the pre-fusion SARS-CoV and SARS-CoV-2 Spikes and shedding of the S1 domain in the latter [30]. Due to its significant overlap with the footprint of 7D6, B9-scFv should likely be considered a 7D6-like RBD-targeting scFv [30]. Overall, B9-scFv binding to this cryptic epitope correlates with its relative cross-reactivity, lack of competition with ACE2 and resistance to RBM mutations found in SARS-CoV-2 VOC.

Despite its modest initial affinity for SARS-CoV-2, it may be possible to improve the binding of an isolated bovine scFv, either by directing AID-mediated mutagenesis, using CRISPR-x [37], or through other diversification methods, such as error-prone PCR or structure-guided affinity maturation [38]. Indeed, all mutagenesis efforts could be entirely focussed on the region encoding the knob domain. It is also probable that with larger initial libraries, binders against more diverse, and cross-reactive epitopes will be isolated. At least some of these would be expected to have a high initial affinity and require fewer mutations, if any, to achieve strong target binding. Previous efforts to select and engineer human antibodies for increased breadth and potency against SARS-like viruses have been remarkably successful [14]. It should be noted, however, that bovine ultralong CDRH3s are unlikely to be amenable to typical oligonucleotide-based CDR diversification methods, due to the length and unique structural requirements of the ultralong CDRH3 [25, 26]. To ensure that improvements in an scFvs potency are not to the detriment of its breadth, any mutagenesis and selection efforts would need to use multiple related antigens for screening.

Clearly, identifying a novel paratope is just the first step in developing a treatment against a new pathogen, as a non-immunogenic scaffold with which to deliver the neutralising agent is crucial. Fortunately, various possibilities exist. Firstly, bovine paratopes have been successfully transferred to a human antibody scaffold with minimal loss of activity [20], although optimisation of a scaffold may be required to achieve stability and good manufacturability [23]. Other options include scFv-Fc fusions [39] or PEGylated scFvs [40], both of which have significantly improved pharmacokinetic profiles over standard scFvs. Nonetheless, a risk remains that the bovine ultralong CDRH3s will be immunogenic when administered intravenously. This has not proven to be a problem for llama paratopes but these CDRH3s are considerably shorter (median 16 aa versus >50 aa) than their bovine counterparts. If this does prove to be an issue for the ultralong bovine CDRH3s, it may be feasible to nebulize modified bovine scFvs, or smaller neutralising fragments such as knob peptides [41], for delivery to the sites of virus entry [42], as has been suggested for heavy chain only (V_H_H) nanobodies.

Pandemic preparedness and responsiveness hinges on the rapid identification of neutralising epitopes. The emergence of three beta-coronaviruses of pandemic potential in the last 20 years indicates the huge risk posed by these viruses and their high incidence of zoonotic transfers. Indeed, with over 50 known SARS-like viruses circulating in bats alone, this will not be the last emergence. Therefore, the generation of libraries of bNAbs that recognise this virus group will allow rapid screening against emerging related viruses. Indeed, even if the antibody has a relatively weak affinity to one virus sub-type, it may be useful against a related pathogen. With a significantly expanded ultralong CDRH3 library, our pipeline has the potential to identify ultralong neutralising antibodies that can be rapidly deployed against new pathogens, without the need for animal immunizations.

## Methods

### Mammalian cell culture

293T cells were grown in Dulbecco’s modified essential medium (DMEM) supplemented with 10 % foetal calf serum, 4 mM L-glutamine, 50 U/ml penicillin and 50 μg/ml streptomycin. Cells were grown in a humified incubator at 37 **°**C with 5% CO_2_.

### Generation of an ultralong scFv library for mammalian cell surface display

To generate the pBovShow expression vector, DNA encoding an scFv expression cassette, with Ig kappa (*IGK*) leader sequence, (GGGS)_x3_ linker, Myc epitope tag and platelet derived growth factor receptor transmembrane domain (PDGFR-TM), was synthesized by IDT (Coralville, Iowa) and cloned into the EcoRV and XbaI sites of the pCS2-MT+ vector (Addgene plasmid #2296). Sequences encoding V_λ_-light chains (LCs) were amplified from bovine whole blood genomic DNA using the primers described [43]. Individual clones were Sanger sequenced and screened for homology to V_λ_-LCs that are known to productively pair with ultralong heavy chains [43]. A sequence with 99% homology to the BLV1H12 V_λ_-LC was cloned into the XhoI and XmaI sites in the scFv expression cassette, downstream of the (GGGS)_x3_ linker. An ultralong heavy chain library was generated using nested PCR and bovine whole blood genomic DNA. First-round amplification was performed with a forward primer hybridising to a unique region upstream of V_1-7_ (5’-GGACCCTCCTCTTTGTGCTCTCAG-3’), whereas the second-round forward primer was V_1_-specific (5’-TCAC<u>GCTAGC</u>CAGGTGCAGCTGCGGGAGTCG-3’; both PCRs used a J_2-4_ specific reverse primer (5’-GGAT<u>AGATCT</u>CTGAGGAGACGGTGACCAGGAG-3’). The final amplicons, flanked by XbaI and BglII sites, were gel purified and cloned into our display vector upstream of the (GGGS)_x3_ linker. The entire ligation reaction was transformed into DH5α competent *E. coli* cells and used to inoculate an overnight midi culture for preparation of polyclonal plasmid DNA encoding the scFv library.

### Purification of His-tagged proteins by immobilised metal affinity chromatography

Full length trimeric Spike protein, residues 1-1208, was kindly provided by the Oxford protein production facility. It has proline substitutions at residues 986 and 987, a GSAS substitution at the furin cleavage site (residues 682-685), a C-terminal T4 fibritin trimerisation motif, an HRV3C protease cleavage site, a TwinStrepTag and an 8xHisTag. It was expressed in mammalian FreeStyle293F cells and purified via immobilised metal affinity chromatography (IMAC). An expression vector for the secretion and purification of His-tagged proteins from mammalian cells was generated by insertion of DNA encoding an IGK leader and 8xHis tag into pCS2-MT+. A DNA fragment encoding the SARS-CoV-2 RBD (aa 319-591) was amplified from pCAGGS-SARS-CoV-2-Spike vector (a kind gift from Keith Grehan) and cloned in frame with the N-terminal IGK leader sequence and C-terminal 8xHis tag using NheI and XhoI sites. DNA fragments encoding the SARS-CoV RBD (aa 319-591) and MERS-CoV RBD (aa 368-586) were synthesised by IDT (Coralville, Iowa) and cloned into the vector using the same restriction sites. All DNA sequences encoding scFvs for purification were sub-cloned from the mammalian display vector into this purification vector using EcoRV and XhoI sites. Proteins were recovered from the supernatants of 293T cultures following transient transfection. Briefly, 5 × 10^6^ 293T cells were transfected with 15 μg of relevant protein expression plasmid in 15 cm^2^ dishes using polyethylenimine (PEI) at a 1:3 DNA:PEI ratio. Complete media was replaced with serum-free media 24 hours after transfection. After a further 96 hours, the supernatant was collected and cleared at 4000 x *g* for 5 minutes before being filtered through a 0.45 μm syringe filter (Fisher). Imidazole (Merck) was added to the cleared supernatants to a final concentration of 10 mM and the supernatants were incubated with 1 ml Nickel resin (Generon; 50% slurry equilibrated in PBS) on a roller at 4 **°**C for 30 minutes to bind His-tagged proteins. The resin was loaded onto a 20 ml econo-column (BioRad) and extensively washed with increasing concentrations of imidazole in PBS (10, 20 and 30 mM). Bound proteins were eluted with 250 mM imidazole in PBS. Protein containing fractions were determined by A^280^ measurements and SDS-PAGE. The required fractions were extensively buffer exchanged into PBS+10% glycerol using a 10 kDa MWCO centrifugal filter (Millipore) and concentrated to >1 mg/ml for storage at -80 **°**C.

### Flow cytometry analysis of cell surface displayed scFv interactions

A standardised staining protocol was followed prior to fluorescence analysis on a Cytoflex S cell analyser (Beckman) and fluorescence-activated cell sorting (FACS) on a FACSMelody (Becton Dickinson; BD). Briefly, 293T cells expressing scFv on the cell surface were detached with trypsin-EDTA (Thermo Fisher Scientific) and washed twice in prechilled sort buffer (1% FCS, 25 mM HEPES-KOH pH 7.9, 1 mM EDTA in PBS). Cells were resuspended in sort buffer to 1 × 10^7^/ml and incubated with the indicated concentrations of target His-tagged proteins for 1 hour at 4 **°**C. After binding, samples were washed twice in sort buffer and incubated with a 1:100 dilution of α-Myc-FITC (Abcam, #Ab1263) and α-His-PE (Abcam, #Ab72467) antibody at room temperature for 10 minutes. Following staining, cells were washed twice in sort buffer and resuspended at 1 × 10^6^/ml for flow cytometry.

### Plasmid recovery

Cells expressing Spike-binding scFvs were purified by flow cytometry, followed by centrifugation at 600 x *g* for 3 min. The respective plasmid expression vectors were recovered by resuspending the pelleted cells in 100 μl of Hirt I solution (0.6% SDS, 10 mM Tris-HCl pH 8.0, 1 mM EDTA; Hirt, 1967) and incubation at room temperature for 10 minutes. Next, 50 μl of Hirt II solution was added (5 M NaCl, 10 mM Tris-HCl pH 8.0, 1 mM EDTA), lysates were mixed and incubated at 4 **°**C overnight. Lysates were centrifuged at 16,000 x *g* for 40 minutes and plasmid DNA was recovered by phenol:chloroform extraction of the supernatant, followed by ethanol precipitation and resuspension in 10 μl of ddH_2_O. DH5α competent *E. coli* (High Efficiency; NEB) were chemically transformed with 5 μl of the recovered plasmid library and incubated for 1 hour at 37 **°**C with shaking at 220 rpm. This was used to inoculate an overnight culture for midi-scale preparation of plasmid scFv library DNA. Three rounds of plasmid-based selection were performed and at each stage Spike binding was verified by transient transfection, while the sequences recovered were characterised by amplicon sequencing.

### Amplicon Sequencing and library size estimation

The scFv library was subjected to amplicon sequencing following rounds 0, 2 and 3 of the plasmid-based enrichment. Briefly, 10 μg of plasmid scFv library from the relevant round of enrichment was digested with EcoRV and BglII to yield a 500 bp fragment spanning the entire ultralong V_H_ sequence. The DNA fragment was gel purified and its concentration adjusted to 20 ng/μl. The resulting fragment library was sequenced by the Illumina-based Genewiz, Amplicon-EZ service to generate 2 × 250 bp paired-end reads. Only reverse reads were used for analysis as they span the whole CDRH3 region and allow accurate characterisation of unique CDRH3 sequences. Raw FastQ files from Genewiz were filtered for quality and converted to Fasta format. Mixcr ([44] https://github.com/milaboratory/mixcr) was used to align the reads to bovine V, D and J segments using *Bos taurus* IMGT libraries for assignment. Unique clonotypes were then assembled and ranked by proportion. The number of unique clonotypes in each case was eventually used to approximate the initial library diversity using the capture-mark-recapture formula 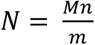, where *N* = heavy chain library size to be estimated, *M* = unique heavy chain sequences recovered in Round 0, *n* = unique heavy chain sequences recovered in Round 2 or Round 3 and *m* = sequences found both in Round 0 and Round 2 or 3.

### Lentivirus particle generation, transduction, and scFv isolation

A lentiviral plasmid for the stable expression of ultralong scFvs was generated by modifying LentiCRISPR v2 (Addgene plasmid #52961, a gift from Feng Zhang). An internal ribosomal entry site (IRES) was cloned downstream of the PDGFR-TM sequence in pBovShow and the whole cassette was inserted between the EF1α core promoter and PuroR gene of LentiCRISPR v2 to generate Lenti-BovShow-IRES-PuroR. The round 3 enriched ultralong scFv library was transferred from pBovShow into this lentiviral vector and lentiviruses were generated by transient transfection of 293T cells. Briefly, 293T cells were seeded at 3 × 10^6^ cells per 10 cm^2^ dish. The next day, 4 μg Lenti-BovShow-IRES-PuroR, 4 μg of pCMVR8.74 packaging vector (Addgene plasmid #22036) and 2 μg of pMD2.G coat protein vector (Addgene plasmid #12259; both gifts from Didier Trono) were mixed with PEI at a 1:3 molar ratio and added to the 10 cm^2^ dish. The medium was changed after 24 hours and lentivirus-containing supernatants were collected at 48- and 72-hours post-transfection. Transduction of 293T cells was achieved by seeding the cells at 30-40% confluency in a 75 cm^2^ flask. The following day, medium was replaced with 9 ml of complete DMEM, plus 1 ml of lentiviral supernatant and polybrene at a final concentration of 5 μg/ml. Medium was replaced after 48 hours with complete medium containing 2 μg/ml puromycin dihydrochloride. After 1 week, puromycin-selected cells were incubated with SARS-CoV-2 Spike (40 nM) and stained with α-Myc-FITC and α-His-PE antibodies as described. Single-cells were purified from the population of Spike-binding cells by flow cytometry using a FACSMelody (BD). Single-cell clones were then assessed for binding to His-tagged SARS-CoV-2 Spike using a Cytoflex S (Beckman) cell analyser. The scFv sequences were amplified from 100 ng of genomic DNA (gDNA) using a forward primer hybridising to the scFv-leader sequence (5’-GACTTT<u>GATATC</u>ATGGAGACAGACACACTCCTG-3’), a J_2-4_ reverse primer 5’-(GGAT<u>AGATCT</u>CTGAGGAGACGGTGACCAGGAG-3’) and HERC II polymerase (Agilent), following the manufacturer’s recommended reaction conditions. Amplicons were analysed by Sanger sequencing and cloned into pBovShow expression and purification vectors.

### scFv binding to Spike variants expressed on the cell surface

293T cells were plated at 0.2 × 10^6^ in 6-well plates 24 hours prior to transfection. The next day, 1 μg of plasmid vector encoding full length SARS-CoV Spike (pCAGGS-SARS-CoV-Spike_Urbani), SARS-CoV-2 Spike (NR-52514 SARS-CoV-2 Spike glycoprotein) or SARS-CoV-2 variant Spike was transfected at a 1:3 DNA to PEI ratio. After 48 hours, cells were detached, washed twice in sort buffer, and incubated with B9-scFv-8xHis at the concentrations indicated. After 1 hour, cells were washed twice in sort buffer, incubated with a 1:100 dilution of α-His-PE antibody (Abcam, #Ab72467) for 10 minutes and washed twice more. Stained cells were resuspended in sort buffer at 1 × 10^6^/ml for fluorescence analysis on a Cytoflex S cell analyser (Beckman). SARS-CoV-2 Spike variants were generated by site-directed mutagenesis with Q5 polymerase (New England Biolabs) and confirmed by Sanger sequencing. Spike cell surface expressions were confirmed by staining with 300 nM of a positive control scFv (CR3022-scFv) that was purified in-house.

### Pseudotype Neutralisation assays

Pseudotyped lentiviral particles were generated by transfecting 3 × 10^6^ 293T cells in a 10 cm^2^ dish with a lentivirus backbone plasmid encoding a luciferase reporter gene (BEI: NR-52516 pHAGE-CMV-Luc2-IRES-ZsGreen-W), a plasmid encoding either the SARS-CoV Spike (pCAGGS-SARS-CoV-Spike_Urbani), SARS-CoV-2 Spike (BEI: NR-52514) or VSV glycoprotein (VSV-G; pMD2.G) and the packaging vectors HDM-Hgpm2 (BEI: NR-52517), HDM-tat1b (BEI: NR-52518) and pRC-CMV-Rev1b (BEI: NR-52519). DNA was transfected at a 1:3 ratio with PEI. Virus particles were collected 48 and 72 post-transfection and resultant lentiviruses serially diluted 1:3 (SARS-CoV and SARS-CoV-2) or 1:10 (VSV-G) on 2.5 × 10^5^ hACE2 expressing 293T cells. In each case, the transduction units per ml (TU/ml) was calculated from the percentage of green cells 48 hours post-transduction, with TU/ml = (number of cells transduced x % positive)/dilution factor. The TU/ml value was calculated from samples with 1-10% green cells. For neutralisation experiments, equal titres of pseudo-typed lentiviruses were incubated with different concentrations of scFv or hACE2-Fc for 1 hour at 37 °C and added to 1.25 × 10^4^ of the ACE2 expressing cells. After 48 hours, the luciferase activity of the infected cells was measured and plotted as a percentage of the virus-only infectivity to generate relative values. All procedures were performed as described by Crawford *et al*., [45] using reagents kindly supplied by BEI, except where indicated. For IC_50_ calculations, SARS-CoV pseudo-typed lentiviruses were incubated with 0.02 - 2 μM CR3022- or 0.04 - 3 μM B9-scFvs and used to transduce 293T-hACE2 cells. The IC_50_ of the scFvs was approximated from the non-linear regression of the summarised log(scFv concentration)-response plots on GraphPad.

### ACE2 competition assay

Initially, His-tagged SARS-CoV RBD (8 nM) was incubated with either ACE2-Fc (300 nM), B9-scFv (5 μM) or CR3022-scFv (5 μM) at 4 **°**C for 30 minutes in sort buffer. Next, aliquots of 0.25 × 10^6^ HEK-293T-hACE2 (BEI: NR-52511) cells were washed in sort buffer and resuspended in one of the pre-incubated RBD samples. After 1 hour at 4 **°**C, cells were washed twice in sort buffer and stained with α-His-PE antibody (1:100). After a final two washes in sort buffer, fluorescence was measured by flow cytometry on a Cytoflex S (Beckman). Overall ACE2 competition was assessed by the reduction in RBD binding to the HEK-293T-hACE2 cell surface as a percentage of RBD-only.

### Hydrogen-deuterium exchange mass spectrometry (HDX-MS)

For HDX-MS experiments, a robot for automated HDX (LEAP Technologies) was coupled to an Acquity M-Class liquid chromatography (LC) system and HDX manager (Waters). Samples comprised protein (SARS-CoV RBD, or SARS-CoV RBD and B9-scFv, at a concentration of 10 µM and 50 µM, respectively) in 50 mM potassium phosphate, pH 8, 0.3 M NaCl. To initiate the HDX experiment, 95 μl of deuterated buffer (50 mM potassium phosphate, pD 8, 0.3 M NaCl) was added to 5 μl of protein-containing solution, and the mixture was incubated at 4 °C for 0.5, 2 and 30 minutes. For each time point and condition, three replicate measurements were performed. The HDX reaction was quenched by adding 100 μl of quench buffer (10 mM potassium phosphate, 0.05 % n-dodecyl-β-D-maltoside (DDM), pH 2.2) to 50 μl of the labelling reaction.

The quenched sample (50 μl) was proteolyzed by flowing through immobilised pepsin and aspergillopepsin columns (Affipro) connected in series (20 °C). The resulting peptides were trapped on a VanGuard Pre-column [Acquity UPLC BEH C18 (1.7 μm, 2.1 mm × 5 mm, Waters)] for 3 min. The peptides were separated using a C18 column (75 μm × 150 mm, Waters, UK) by gradient elution of 0 - 40% (v/v) buffer B in buffer A where buffer A is H_2_O, 0.3% v/v and buffer B is acetonitrile, 0.1% v/v formic acid over 7 min at 40 μl min^−1^.

Peptides were detected using a Synapt G2Si mass spectrometer (Waters) operating in HDMS^E^ mode, with dynamic range extension enabled. Ion mobility (IM) separation was used to separate peptides prior to collision induced dissociation (CID) fragmentation in the transfer cell. CID data were used for peptide identification, and uptake quantification was performed at the peptide level. Data were analysed using ProteinLynx Global Server (PLGS) (v3.0.2) and DynamX (v3.0.0) software (Waters). Search parameters in PLGS were as follows: peptide and fragment tolerances = automatic, minimum fragment ion matches = 1, digest reagent = non-specific, false discovery rate = 4. Restrictions for peptides in DynamX were as follows: minimum intensity = 1000, minimum products per amino acid = 0.3, max sequence length = 25, max ppm error = 5, file threshold = 3. The software Deuteros was used to identify peptides with statistically significant increases/decreases in deuterium uptake and to prepare Wood’s plots. The raw HDX-MS data have been deposited to the ProteomeXchange Consortium via the PRIDE partner repository with the dataset: PXD032965. A summary of the HDX-MS data, as recommended by reported guidelines is shown in Supplementary Table 1.

### Disruption of the B9-scFv knob domain

The sequence encoding B9-scFv was mutated in the BovShow cell surface expression vector; residues _123_YNCRPAVWY_131_ of the B9-scFv knob domain (B9-WT) were replaced with the irrelevant amino acid sequence _123_ETCYYGSGL_131_ by site-directed mutagenesis (B9-Mut) with Q5 polymerase (New England Bioloabs). 293T cells were transfected with either B9-WT or B9-Mut plasmid DNA and cell-surface expressed scFvs were tested for binding to purified SARS-CoV RBD (200 nM) using the standard staining protocol.

### Structural models used in this study

The PDB files used in this study were 4K3D, 6M0J, 5×5B and 6VXX. All structural figures used in this study were generated in UCSF Chimera.

### Statistical information

Flow cytometry experiments include a positive and unstained negative control and were performed at least in triplicate and/or with sufficient replicates to ensure statistically significant data (except Fig. 5c that was performed in duplicate). Quantification of binding is determined using mean fluorescence intensity via CytExpert2.4 and is plotted to show the standard error of the mean. The K_D_ for interactions between cell surface scFvs and recombinant RBD proteins was estimated by non-linear analyses of the log(molarity)-response plots on GraphPad.

Pseudotype neutralisation assays were performed at least in triplicate to calculate the standard error of % neutralisation (compared to the negative control) at each concentration of scFv. IC_50_ values are calculated from the nonlinear regression of log(molarity) of scFv versus % neutralisation.

For differential HDX-MS, peptide-level significance testing was implemented in Deuteros 2.0 to identify peptides with significant differences in deuterium uptake in the bound state. A hybrid significance test was used that first evaluates if the difference in deuterium uptake between two states is greater than a threshold value that corresponds to a significance level of p < 0.01. This was followed by a Welch’s *t*-test to confirm that the differences are significant.

## Data availability

The raw HDX-MS data have been deposited to the ProteomeXchange Consortium via the PRIDE partner repository with the dataset: PXD032965

## Acknowledgements

We gratefully acknowledge funding from the BBSRC (BB/V01384X/1 to JMB, Sinisa Savic and PGS), a MRC studentship (MR/N013840/1 to MJB) and the Wellcome Trust (Joint Investigator Award: 110145 to PGS). ANC acknowledges support of a Sir Henry Dale Fellowship jointly funded by the Wellcome Trust and the Royal Society (Grant Number 220628/Z/20/Z) and a University Academic Fellowship from the University of Leeds. Funding from BBSRC (BB/M012573/1) enabled the purchase of mass spectrometry equipment. We are grateful to the Oxford protein production group for the trimeric SARS-CoV-2 Spike protein and BEI for a series of reagents. Specifically, the following reagents were obtained through BEI Resources, NIAID, NIH: (a) Human Embryonic Kidney cells (HEK-293T) expressing the human angiotensin-converting enzyme 2, HEK-293T-hACE2 cell line, NR-52511 and (b) Modified pαH Vector Containing the Human Angiotensin-Converting Enzyme 2, NR-52565. The following reagents were also obtained through BEI resources and produced under HHSN272201400008C: (a) Plasmid Set for Anti-SARS Coronavirus Human Monoclonal Antibody CR3022, NR-53260 and (b) Vector pCAGGS Containing the SARS-Related Coronavirus 2, Wuhan-Hu-1 Spike Glycoprotein Gene (soluble, stabilized), NR-52394. We also gratefully acknowledge Professor Nicola Stonehouse and Dr. Keith Grehan for the pCAGGS-SARS-CoV-2-Spike and pCAGGS-SARS-CoV-Spike_Urbani expression vector.

## Author contributions

M.J.B., P.G.S., A.N.C. and J.M.B. designed the study. M.J.B., J.N.F.S., T.C.M. and A.N.C. acquired, and analysed the data. M.J.B., P.G.S. and J.M.B. wrote the manuscript. All authors participated in discussion and interpretation of the data, contributed to experimental design and edited the manuscript.

## Materials and correspondence

Requests for materials and manuscript correspondence should be sent to Joan Boyes

