## Supplemental Figures and Table for "An Ultralong Bovine CDRH3 that Targets a Conserved, Cryptic Epitope on SARS-CoV and SARS-CoV-2"

a

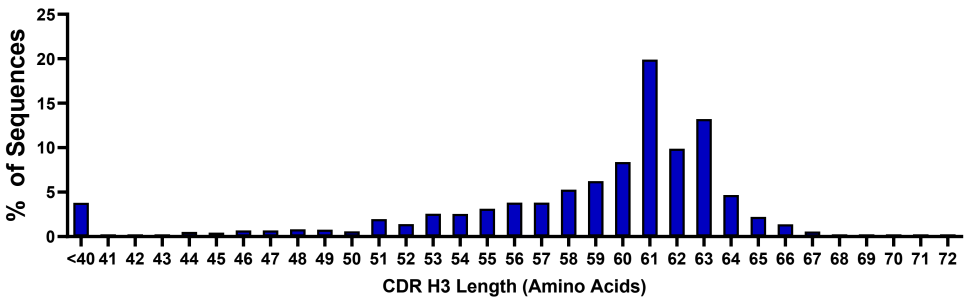


b

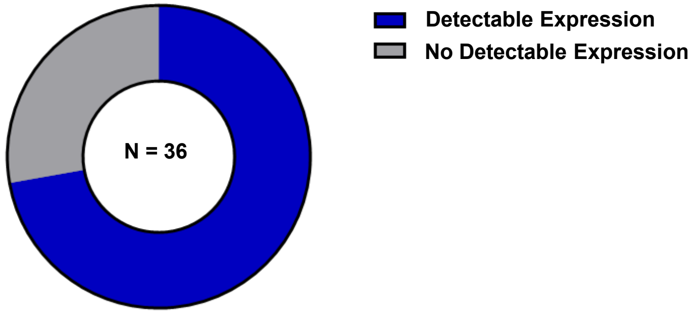


c

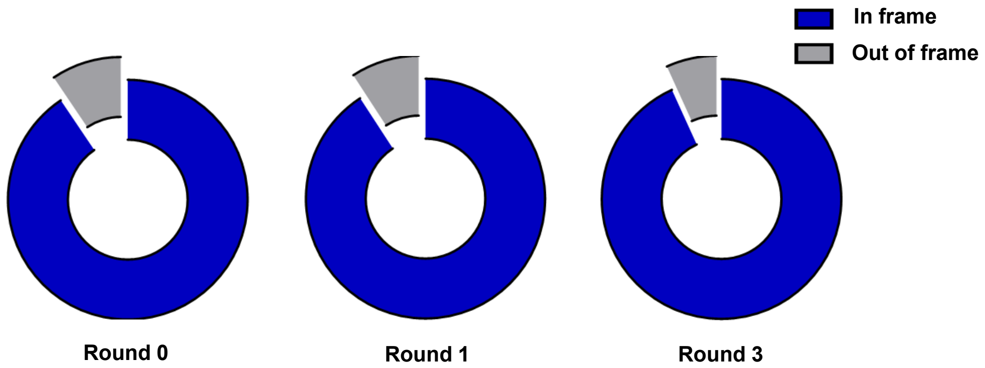


| | Round 0 | Round 2 | Round 3 |
|---|---|---|---|
| **Unique** | 3168 | 379 | 250 |
| **Non-unique** | N/A | 260 | 102 |

| | |
|---|---|
| **Estimated Library Size** | |
| **Round 2** | **Round 3** |
| 4617 | 7764 |

**Supplementary Figure 1**

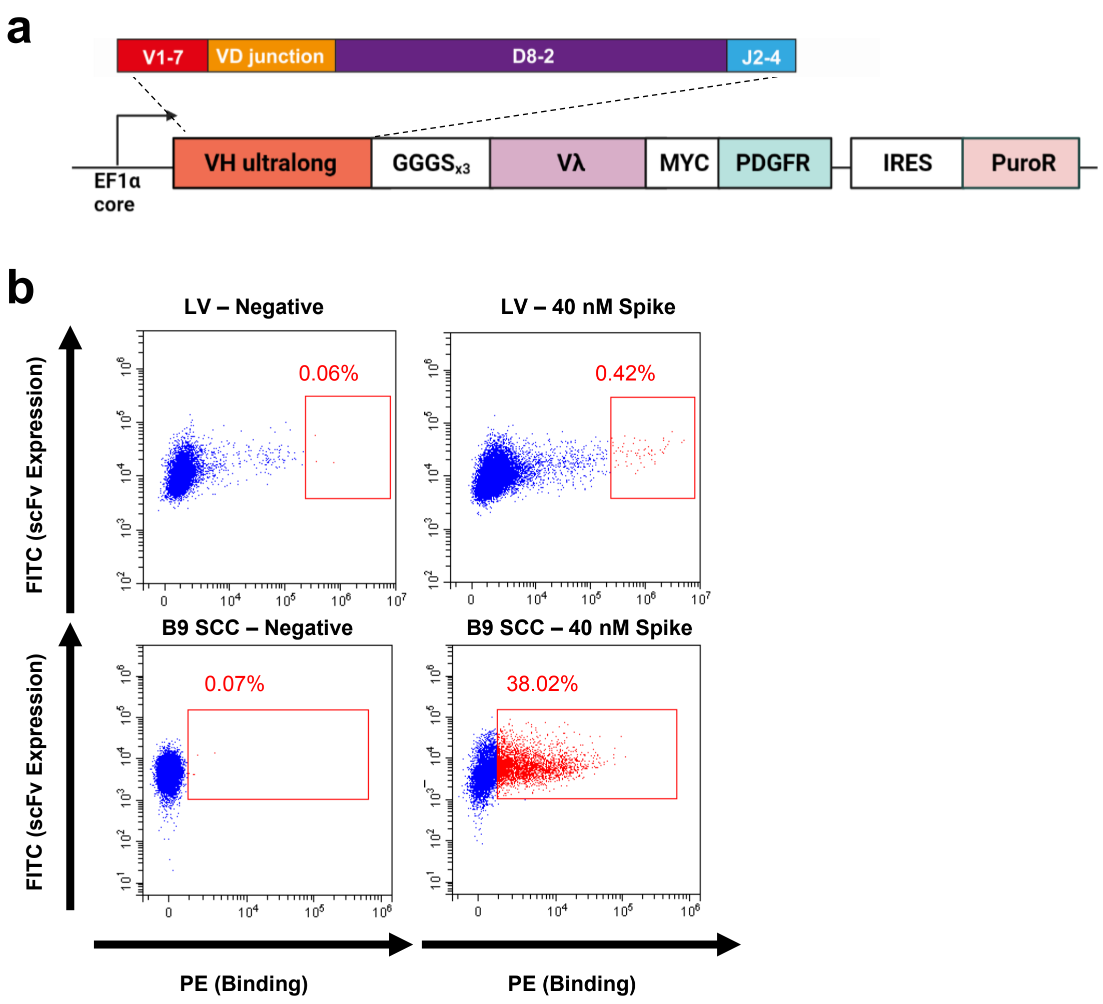


Supplementary Figure 2

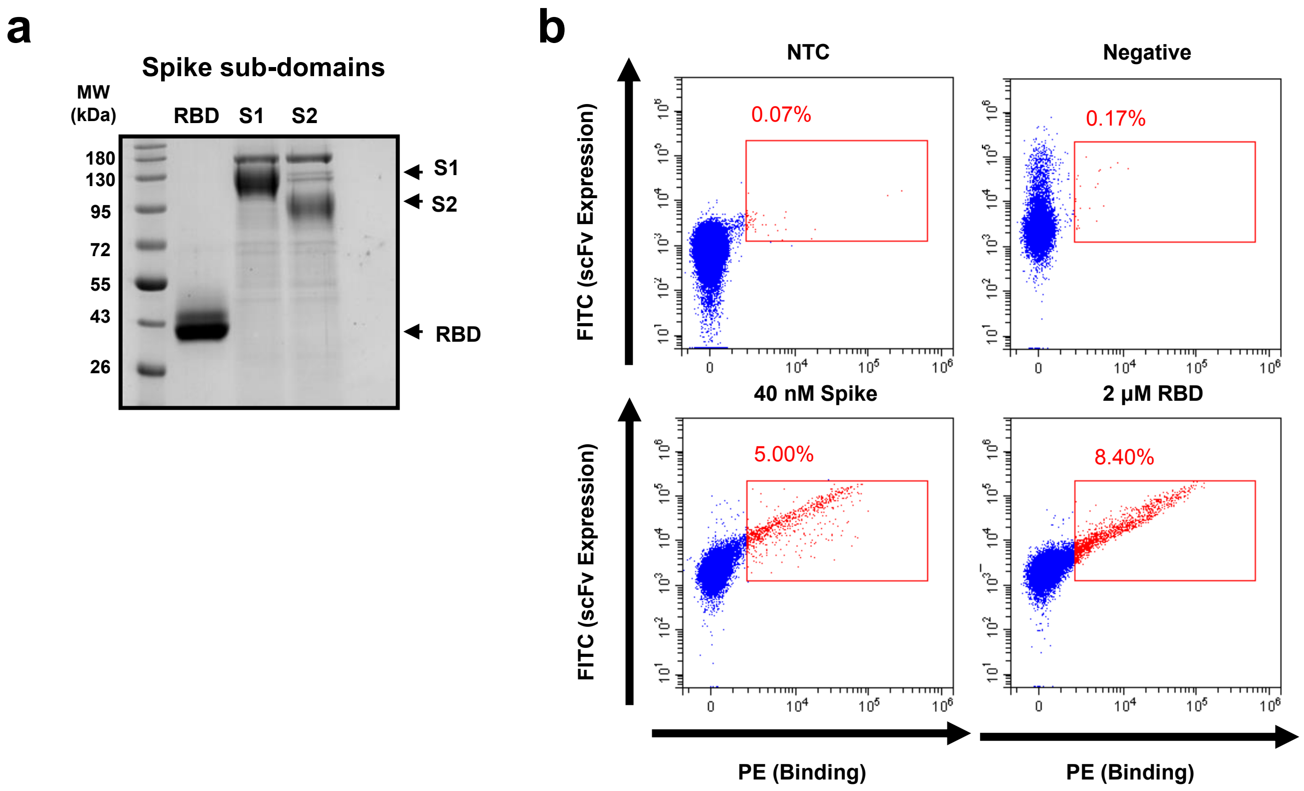


**Supplementary Figure 3**

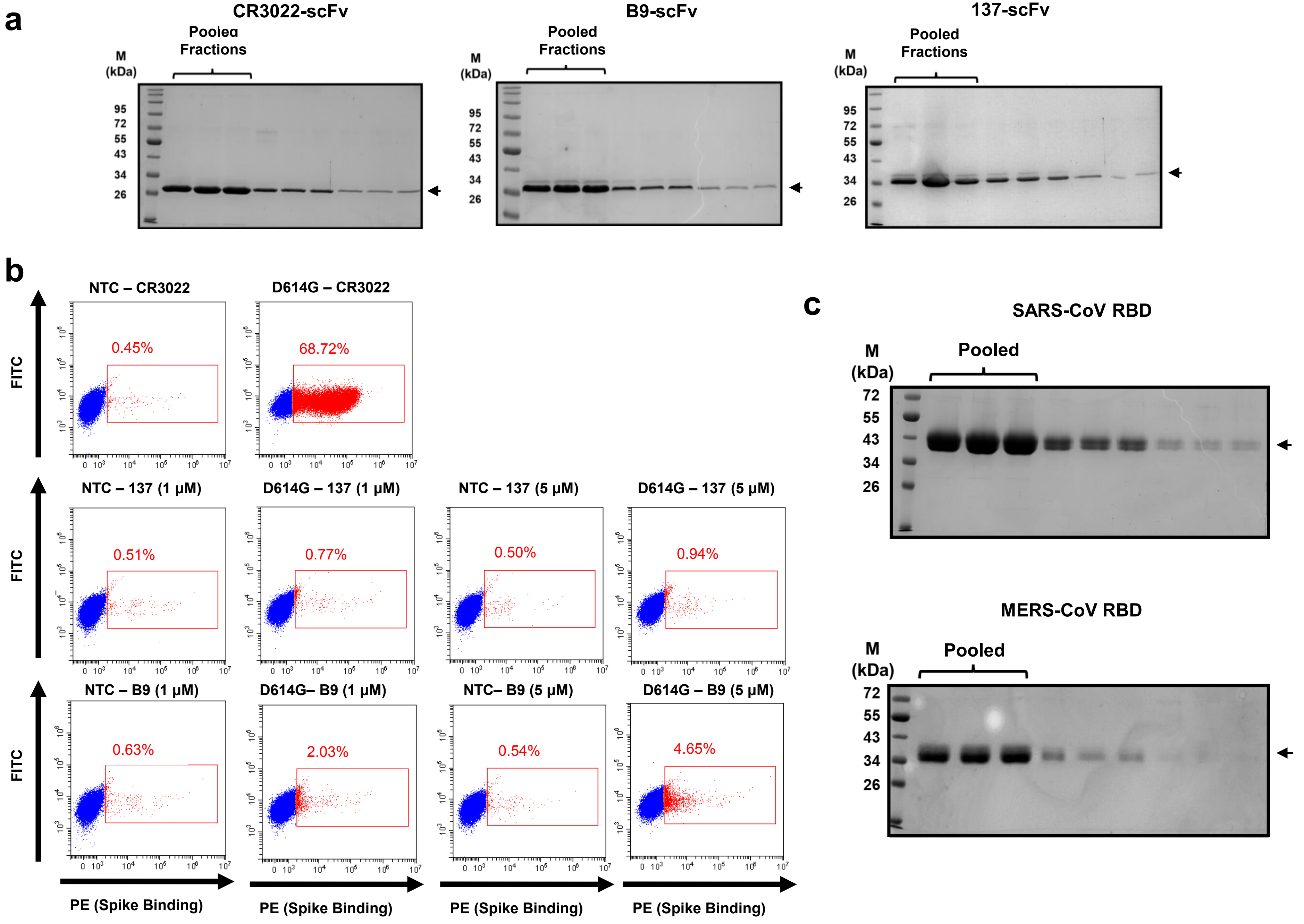


Supplementary Figure 4

**a**

| Protein | Start | End | Peptide | State | Average uptake difference | Standard Deviation |
|---|---|---|---|---|---|---|
| SARS-CoV-RBD | 449 | 467 | YNYKYRYLRHGKLRPFERD | RBD + B9-scFv | -0.6526 | 0.0411 |
| SARS-CoV-RBD | 452 | 467 | YKYRYLRHGKLRPFERD | RBD + B9-scFv | -0.7301 | 0.1911 |
| SARS-CoV-RBD | 456 | 467 | LRHGKLRPFERD | RBD + B9-scFv | -0.5460 | 0.0810 |
| SARS-CoV-RBD | 551 | 565 | VLTPSSKRFQPFQQFGRD | RBD + B9-scFv | -0.3821 | 0.0924 |

**b**

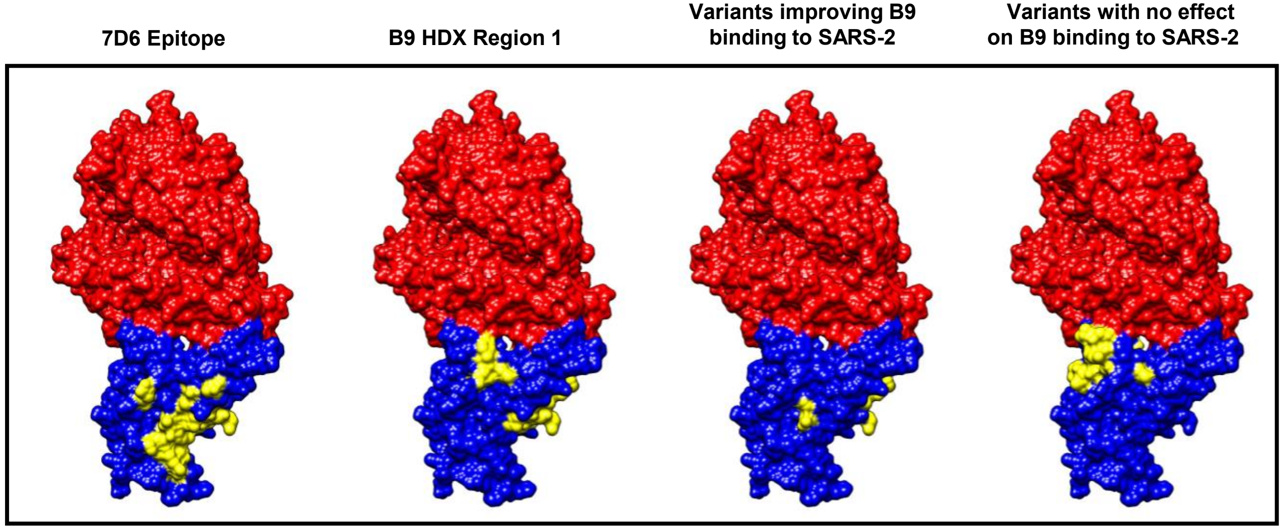


**c**

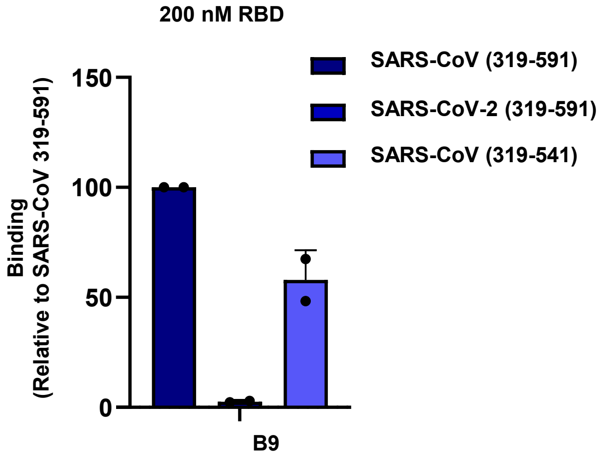


**d**

| Mutations |
|---|
| A348P |
| N354E |
| P384A |
| Mut402-406 (I402V, R403K, E406D) |
| I434L |
| Mut438-447 (S438T, N439R, L441I, S443A, K444T, V445S, G446T) |
| L452K |
| Mut455-462 (L455Y, F456L, K458H, S459G, N460K, K462R) |

**e**

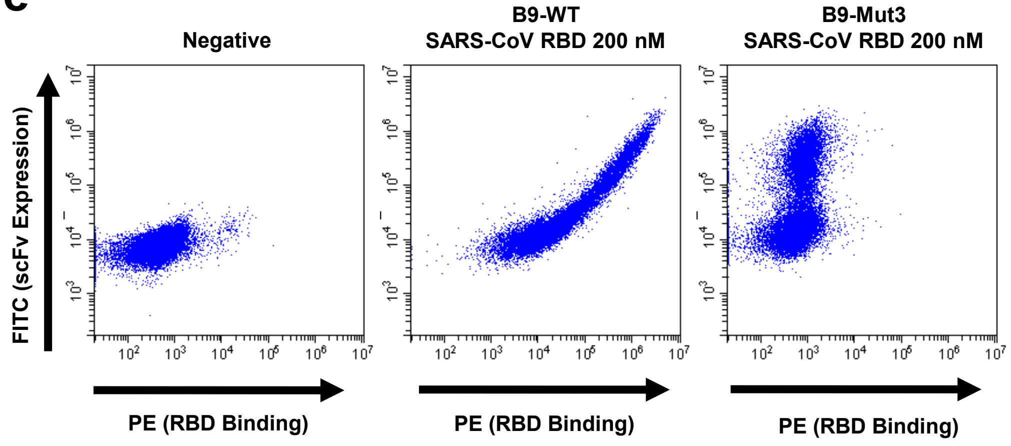


B9-WT CTTVHQRDSCPYDSSHSQYCSYGWSCGVYNCRPAVWYRYGSCGTCDNLFSYVDAW

B9-Mut3 CTTVHQRDSCPYDSSHSQYCSYGWSCGVETCYYGSGLRYGSCGTCDNLFSYVDAW

**Supplementary Figure 5**

**Supplementary Figure 1: Estimation of scFv library diversity. a)** Histogram showing the proportion of heavy chain amino acid sequences with CDRH3 between 40 and 72 amino acids. Heavy chain amino acid sequences were determined from the amplicon sequencing of the Round 0 plasmid scFv library. **b)** Random pBovShow-scFv clones were picked (N=36), transfected into 293Ts and tested for expression by staining with α-Myc-FITC fluorescent antibody after 48 hours. The positive fraction is indicated in blue. **c)** Clonotypes were assembled from amplicon sequencing of our scFv plasmid library at three different time points during the Spike-binding enrichment (Round 0, Round 2 and Round 3). The resultant clonotype assemblies were used to assess the fraction of productive and non-productive heavy chain sequences. The number of unique clonotypes in each case was also used to approximate the initial library diversity using the capture-mark-recapture formula $N = \frac{Mn}{m}$ , where $N$ = heavy chain library size to be estimated, $M$ = unique heavy chain sequences recovered in Round 0, $n$ = unique heavy chain sequences recovered in Round 2 or Round 3 and $m$ = sequences found both in Round 0 and Round 2 or 3. The calculated level of diversity (<1 x $10^4$ unique sequences) is lower than that of typical libraries (5 x $10^7$) and is likely due to the library being generated from leucocyte, rather than spleen, DNA.

**Supplementary Figure 2: Lentivirus transduction of scFv library. a)** Cartoon showing the cloning of the enriched ultralong $V_H$ library into the lentivirus vector used to generate stable cell lines. **b)** Upper: FACS plots showing stable polyclonal 293T cells transduced with enriched lentivirus scFv library incubated without (left) and with (right) 40 nM Spike. Lower: Single cell clone (SCC) B9-scFv incubated without (left) and with (right) 40 nM Spike. The red boxes show cells that express scFvs that bind Spike. All samples were stained with α-His-PE and α-Myc-FITC.

**Supplementary Figure 3: Isolation of an ultralong scFv that binds to the wildtype SARS-CoV-2 Spike and receptor binding domain. a)** SDS-PAGE of purified SARS-CoV-2 Spike sub-domains. **b)** FACS plots showing, Upper: non-transfected 293T cells (left) and 293T cells transfected with B9-scFV (right). Lower: 293T cells transfected with B9-scFv and incubated with 40 nM Spike (left) or 2 μM RBD (right). All samples were stained with α-Myc-FITC and α-His-PE.

**Supplementary Figure 4: Purified B9-scFv binds to Wuhan-Hu-1 (+D614G) SARS-CoV-2 Spike expressed on the surface of 293T cells. a)** SDS-PAGE gels of purified CR3022-, 137- and B9-scFvs. **b)** Upper: Non-transfected (NTC) and 293T cells transfected with Wuhan-Hu-1 (+D614G) Spike gene were incubated with 200 nM CR3022-scFv to confirm cell surface

Spike localization. Middle & lower: Non-transfected (NTC) and 293T cells transfected with an expression vector for SARS-CoV-2 Spike protein were incubated with 1 and 5 µM 137-scFv (middle; negative control) or 1 and 5 µM B9-scFv (lower). All samples were stained with α-His-PE antibody (1:100). **c)** SDS-PAGE of purified SARS-CoV (upper) and MERS-CoV (lower) RBD.

**Supplementary Figure 5: Localising the epitope of B9-scFv to a cryptic site by hydrogen-deuterium exchange and site-directed mutagenesis. a)** Peptides of SARS-CoV RBD protected from deuterium uptake in the presence of B9-scFv following a 2-minute exposure to deuterated buffer. **b)** Surface maps of monomeric SARS-CoV-2 RBD (Blue) bound to hACE2 (red) generated in UCSF Chimera (PDB: 6M0J). Regions of interest are highlighted in yellow. There is significant overlap between the footprint of 7D6 (right) and the proposed epitope of B9. **c**) 293T cells expressing B9-scFv on the cell surface were incubated with 200 nM of the recombinant RBD proteins listed. Binding was measured as the mean fluorescence intensity (MFI) and is presented relative to SARS-CoV (319-591). Error bars represent standard deviation (N=2). **d)** SARS-CoV-2 to SARS-CoV mutations tested in Fig. 5c, d. **e)** The knob sequence of B9-scFv is disrupted by the replacement of eight residues with an irrelevant amino acid sequence (ETYYGSGL). 293T cells were transfected with either B9-scFv-WT or B9-scFv-Mut. 48 hours post-transfection, cells were incubated with 200 nM of SARS-CoV RBD and stained with α-His-PE (1:100).

**Supplementary Table 1: HDX Data Summary Table.** SD = standard deviation, CI = confidence interval.

| Data Set | SARS-CoV RBD | SARS-CoV RBD + B9-scFv |
|---|---|---|
| HDX reaction details | 50 mM potassium phosphate, pD 8, 0.3 M NaCl, 4 ºC | 50 mM potassium phosphate, pD 8, 0.3 M NaCl, 4 ºC |
| HDX time course (min) | 0.5, 2, 30 min | |
| HDX control samples | Maximally labeled controls were not performed. | |
| Back-exchange | ~ 30 % | |
| # of Peptides | 81 | 81 |
| Sequence coverage | 75.67% | 75.67% |
| Average peptide length / Redundancy | 11.04 / 3.94 | 11.04 / 3.94 |
| Replicates (biological or technical) | 3 (technical) | 3 (technical) |
| Repeatability | 0.0476 (average SD) | 0.0465 (average SD) |
| Significant differences in HDX<br>(delta HDX > X D) | Reference | Hybrid Significance test: 99% CI: 0.27Da/ p-value <-0.01 |
